# BackBone Builder (B3): A modular Golden Gate standard with a compatible *Agrobacterium* parts library

**DOI:** 10.64898/2026.02.15.705982

**Authors:** Jonas De Saeger, Mattias Vermeersch, Stijn Aesaert, Laurens Pauwels, Thomas B. Jacobs

**Affiliations:** Department of Plant Biotechnology and Bioinformatics, Ghent University, 9052 Ghent, Belgium; Center for Plant Systems Biology, VIB, 9052 Ghent, Belgium

## Abstract

*Agrobacterium-*mediated transformation relies on binary vectors in which T-DNA and virulence genes are maintained on separate replicons. While Golden Gate cloning has become standard for T-DNA assembly, no modular framework exists for systematic construction of Agrobacterium vector backbones. Here, we present BackBone Builder (B3), a Golden Gate-based platform for combinatorial backbone assembly. B3 uses the Type IIS enzyme PaqCI to minimize domestication and enables one-pot assembly of nine backbone modules plus a selectable cloning cassette. The system is compatible with GreenGate and remains independent of downstream cloning strategies. We generated a library of 42 backbone components, supporting a theoretical design space exceeding 370,000 constructs. A 4 x 4 origin-of-replication (ORI) matrix combining four Escherichia coli and four *Agrobacterium* ORIs assembled with 100% efficiency and functioned robustly in bacterial and plant contexts. Reporter expression reflected expected ORI-dependent patterns in *E. coli, Agrobacterium*, and *Nicotiana benthamiana*. A B3-derived maize transformation backbone achieved stable transformation efficiencies comparable to established vectors. B3 establishes a standardized and extensible framework for rational engineering of *Agrobacterium* binary vector architecture.

## Dear Editor

The ability of *Agrobacterium tumefaciens* and related Rhizobiales to transfer DNA into host genomes is foundational to plant biotechnology. This process is enabled through the binary vector system, in which the T-DNA and virulence (*vir*) genes reside on separate replicons. In the most widely used configuration, the large native Ti plasmid is split into a helper plasmid containing the *vir* region and a compact binary vector used to clone gene(s) of interest (De Saeger et al., 2021). While early binary vectors contained only a broad-host-range origin of replication (ORI), a bacterial selectable marker, and the T-DNA (Bevan, 1984), modern backbones now provide a variety of alternative designs and optional modules (De Saeger et al., 2021). These include dual-T-DNA systems (Komari et al., 1996), alternative ORIs for *Agrobacterium* and *E. coli* (Hamilton, 1997), the addition of toxic genes behind the left border to avoid regenerating plants containing vector backbone (Hanson et al., 1999), and modified T-DNA borders (Kuraya et al., 2004; Sardesai et al., 2018). Despite the widespread use of Golden Gate cloning for T-DNA assembly (Lampropoulos et al., 2013), no standardized framework exists for the modular construction of *Agrobacterium* vector backbones. To fill this gap, we introduce BackBone Builder (B3), a Golden Gate-based system for flexible backbone assembly.

To develop the B3 cloning standard, we identified several key requirements. First, the system needed to assemble multiple parts in a single reaction to facilitate throughput. Second, part domestication had to be minimized to avoid unnecessary sequence changes, which is especially important for regulatory regions and ORIs. Third, the system had to remain compatible with existing downstream cloning methods. Given these requirements, we adopted a Golden Gate scheme using the restriction enzyme PaqCI (isoschizomer of AarI), whose seven-base-pair recognition site greatly reduces the need for domestication. We designed a high-fidelity overhang set (Supplementary Data S1) consisting of 10 overhang pairs that enable assembly of nine backbone modules plus a cloning cassette (containing *ccdB* in the design presented here). The B3 framework is also compatible with the GreenGate standard, which uses the restriction enzyme BsaI (Lampropoulos et al., 2013). Empty B3 entry vectors contain GreenGate A-G sites, allowing B3 to generate new *Agrobacterium* vector parts using either Golden Gate or Gibson assembly. Depending on the cloning cassette incorporated during assembly, destination vectors compatible with any cloning system can be generated (Figure 1).

**Figure 1:**
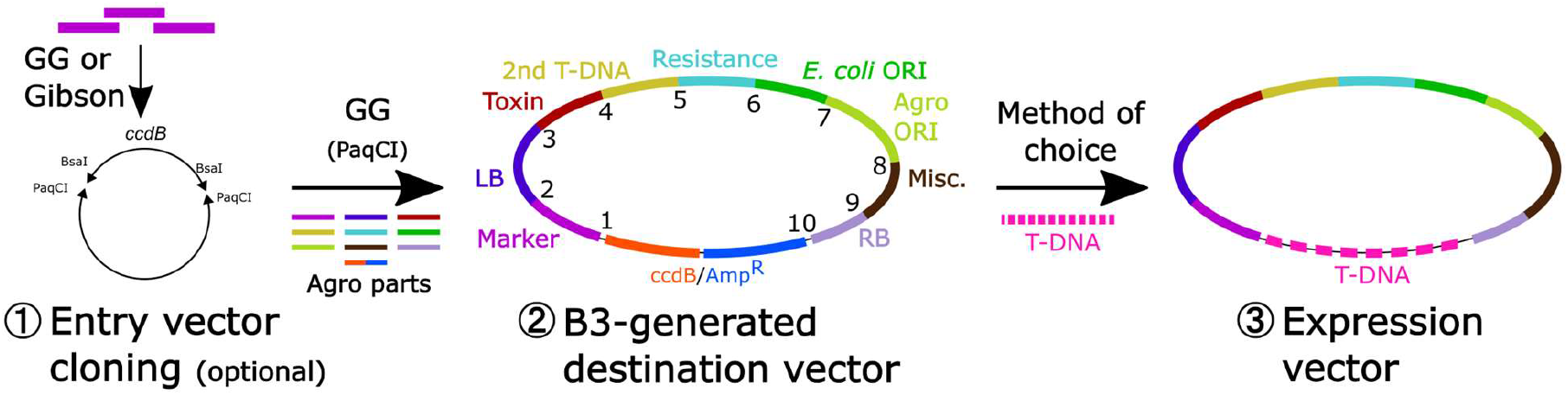
The BackBone Builder (B3) cloning scheme: (1) Entry vector cloning: Inserts can be cloned into empty entry vectors using Golden Gate (via BsaI sites following the GreenGate standard) or Gibson Assembly. Alternatively, new entry clones can be synthesized *de novo*. This step is only required when new parts need to be generated. **(2) B3-generated destination vector:** The B3 system assembles all backbone parts into a complete destination vector using the Type IIS restriction enzyme PaqCI. Each entry plasmid is flanked by PaqCI sites with unique overhangs that determine its position in the final construct. These entry plasmids contain the individual backbone modules, including the ORI, bacterial selectable marker, and other required elements. In the arrow leading into step 2, only the individual backbone fragments are shown; the plasmid backbone is omitted for clarity. **(3) Expression vector:** The final expression construct is generated by inserting the desired T-DNA cargo into the B3 destination vector using a cloning method of choice, depending on the cloning cassette used during destination vector assembly. **Abbreviations:** GG, Golden Gate; LB, Left Border; RB, Right Border; T-DNA, transfer DNA; ORI, origin of replication; AmpR, ampicillin resistance; ccdB, control of cell death B.

We generated the full set of empty B3 entry plasmids and generated a collection of *Agrobacterium* backbone parts (Supplementary Table S1) via Golden Gate or Gibson assembly (Supplementary Methods). The *Agrobacterium* parts kit currently contains 42 components that can be readily combined to assemble new backbone designs, supporting a design space of >370,000 constructs. Additional parts can be cloned into the empty entry plasmids as needed. To evaluate the system’s performance, we constructed a set of novel destination vectors using the GreenGate standard, keeping all parts constant except for the ORIs. The value of having flexibility in the *Agrobacterium* ORI has recently been highlighted through directed evolution approaches aimed at improving transformation efficiency (Szarzanowicz et al., 2025). We tested four *E. coli* ORIs (pUC, ColE1, p15A, and Phage P1) and four *Agrobacterium* ORIs (WKS1, pVS1, OriV, and pRI). These sixteen combinations yielded cloning efficiencies of 100%, as all 80 screened colonies (5 per assembly) were correctly assembled as shown by restriction digest and full plasmid sequencing of one clone per assembly (Supplementary Data S2 and S3). To evaluate whether each construct functioned correctly in both bacterial and plant contexts, we assembled the visual reporters pJ23119::eforCP (Alieva et al., 2008) together with pPcUBI::RUBY (He et al., 2020) into these 16 destination vectors and transformed the verified plasmids into *Agrobacterium* LBA4404 for leaf infiltration assays (Figure 2A). Expression of eforCP in *E. coli* (Figure 2B) followed the expected ORI copy-number–dependent patterns (Supplementary Table S1). In *Agrobacterium* fewer quantitative copy-number studies are available, but our results are consistent with the anticipated ranking of pVS1 > oriV > pRI (Figure 2C). The WKS1 ORI is a recent addition to the *Agrobacterium* toolbox and has not yet been characterized for plasmid copy number; however, in our experiments it produced the highest eforCP expression levels (Kocaoglan et al., 2024). We also observed an interaction effect involving the Phage P1 ORI, which reduced expression when combined with WKS1 (Figure 2C). Finally, transient expression of RUBY in *Nicotiana benthamiana* (Figure 2D) largely mirrored the pattern observed in *Agrobacterium*, consistent with the well-established link between *Agrobacterium* plasmid copy number and *in planta* transient expression output (Szarzanowicz et al., 2024).

**Figure 2:**
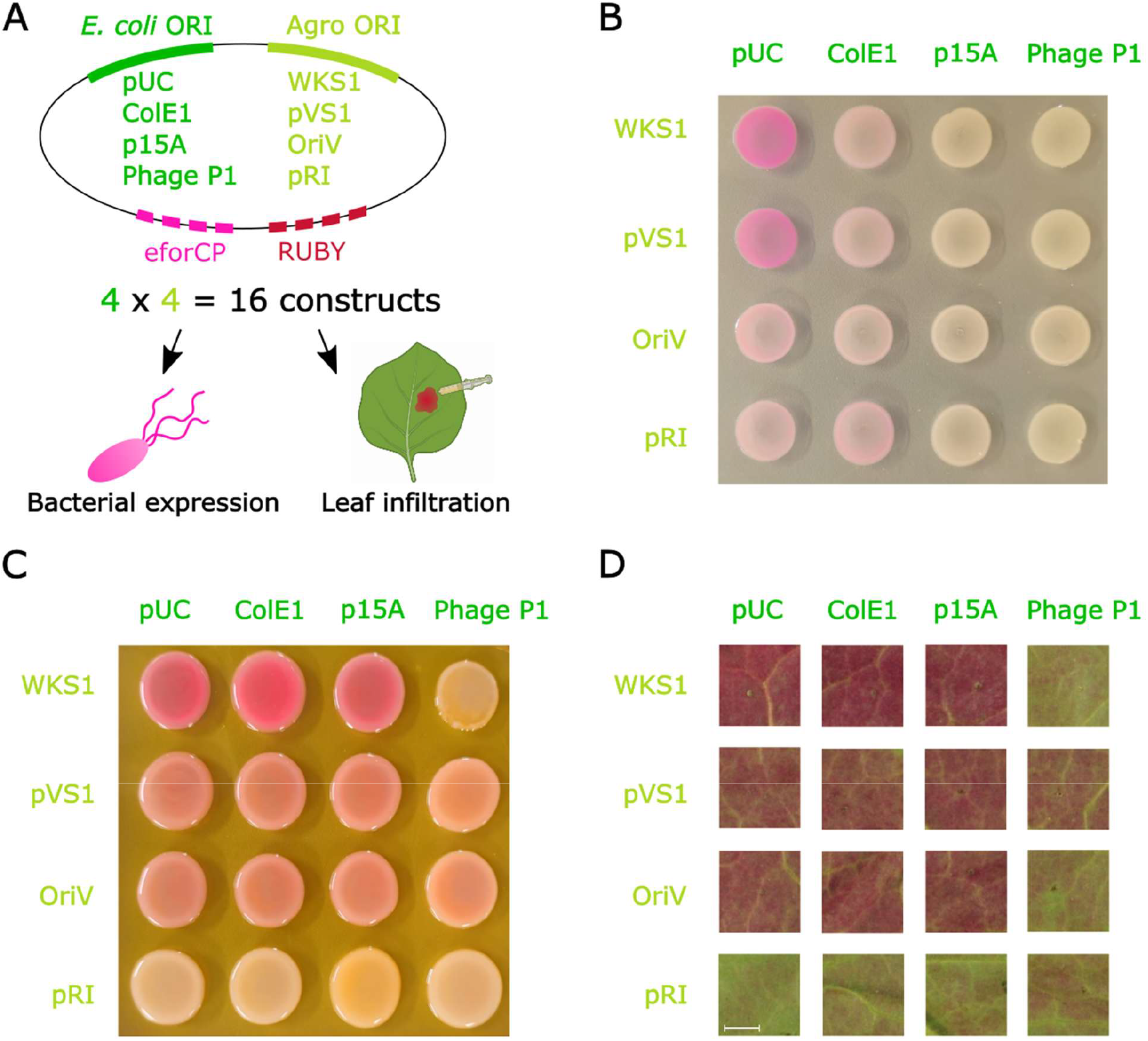
Performance of B3-generated vectors in bacteria and plants. **(A)** Schematic overview of the 4 × 4 origin-of-replication (ORI) matrix. Four *E. coli* ORIs (pUC, ColE1, p15A, Phage P1) were combined with four *Agrobacterium* ORIs (WKS1, pVS1, OriV, and pRI), generating 16 destination vectors. pPcUBI::RUBY and pJ23119::eforCP were assembled into each destination vector, transformed into *E. coli, Agrobacterium tumefaciens* LBA4404, and agroinfiltrated into *Nicotiana benthamiana*. **(B)** *E. coli* expression of eforCP across all 16 ORI combinations. **(C)** *Agrobacterium* expression of eforCP across all 16 ORI combinations. Fluorescence images corresponding to panels B and C are shown in Supplementary Figure S5. **(D)** Transient RUBY expression in *N. benthamiana* leaves following agroinfiltration with each construct at 5 days post-infiltration (6 dpi shown in Supplementary Figure S6). Scale bar = 5 mm.

To evaluate stable plant transformation, we used B3 to assemble the destination vector pB3G3HI-AG, which is based on the maize transformation backbone pG3HI-AG. Minor modifications were introduced, including the removal of nonessential sequences. Another difference is the cloning cassette (either *ccd*B or sfGFP), which is used only during assembly and is not retained in the final expression construct (Vandeputte et al., 2024; Supplementary Data S7). We generated an expression vector with pB3G3HI-AG using GreenGate cloning by introducing pZm13::GFP (Hamilton et al., 1992), together with the transformation booster sequence (TBS) (Hily et al., 2009). A total of 454 embryos from eight different ears from the B104 inbred were transformed, yielding 24 hygromycin-resistant plantlets derived from 17 embryos (3.7%), a transformation efficiency consistent with our routine performance using pG3HI-AG (Vandeputte et al., 2024). Event quality was assessed by digital PCR, identifying seven single-copy, backbone-free plants derived from five explants (22.6%), again in line with previous results (Supplementary Data S8). These data demonstrate that the pB3G3HI-AG backbone is fully functional for stable maize transformation.

In summary, B3 establishes a modular and standardized framework for assembling *Agrobacterium* binary vector backbones, while remaining agnostic with respect to downstream cloning protocols. Although this work focuses on binary *Agrobacterium* vectors, the B3 standard can be readily extended to other complex systems such as super-binary or ternary vectors (De Saeger et al., 2021), yeast shuttle vectors (Gnügge and Rudolf, 2017), or viral delivery constructs (Mahmood et al., 2023).

