## Supplemental Information for "BackBone Builder (B3): A modular Golden Gate standard with a compatible *Agrobacterium* parts library"

**SUPPLEMENTARY INFORMATION OVERVIEW**

| ITEM | DESCRIPTION | FORMAT |
|---|---|---|
| Supplementary Methods | Detailed overview of B3 scheme and experimental procedures. | Supplementary PDF |
| Table S1 | Elements in the BackBone Builder (B3) cloning scheme. | Supplementary PDF |
| Table S2 | Overview of all empty B3 entry plasmids and *Agrobacterium* parts library. | Supplementary PDF |
| Table S3 | List of primers used in this study | Excel (.xlsx) |
| Table S4 | List of plasmids used in this study | Excel (.xlsx) |
| Table S5 | Overview of B3 assemblies and corresponding expected restriction digestion patterns. | Supplementary PDF |
| Figure S1 | The BackBone Builder (B3) cloning scheme | Supplementary PDF |
| Figure S2 | Expected digestion patterns for AG1-AG4 | Supplementary PDF |
| Figure S3 | Expected digestion pattern for AG5-AG8 | Supplementary PDF |
| Figure S4 | Expected digestion pattern for AG9-AG16 | Supplementary PDF |
| Figure S5 | Fluorescence images corresponding to Figure 2B (*E. coli*) and Figure 2C (*Agrobacterium*) eforCP expression. | Supplementary PDF |
| Figure S6 | *Nicotiana benthamiana* leaf infiltration (6 dpi) | Supplementary PDF |
| Figure S7 | Comparison of pG3HI-AG and pB3G3HI-AG plasmids. | Supplementary PDF |
| Data S1 | High-Fidelity set of PaqCI overhangs. | Supplementary PDF |
| Data S2 | B3 cloning results (diagnostic digests) | Supplementary PDF |
| Data S3 | Verification of 16 *E. coli* – *Agrobacterium* combinations. Contains in-silico designed and WPS verified maps for comparison. | Folder (.gb files) |
| Data S4 | B3 empty entry clone plasmid maps (E1-E9) | Folder (.gb files) |
| Data S5 | *Agrobacterium* parts plasmid maps (A1-A42) | Folder (.gb files) |
| Data S6 | 16 ORI combination plasmid maps containing eforCP and RUBY cassettes | Folder (.gb files) |
| Data S7 | Plasmid maps for the destination vectors (pG3HI-AG, pB3G3HI-AG) and the maize expression construct pB3G3HI-Zm13GFP-TBS. | Folder (.gb files) |
| Data S8 | pB3G3HI-Zm13GFP-TBS maize transformation efficiency | Excel (.xlsx) |

### SUPPLEMENTARY METHODS

Overview of the BackBone Builder scheme

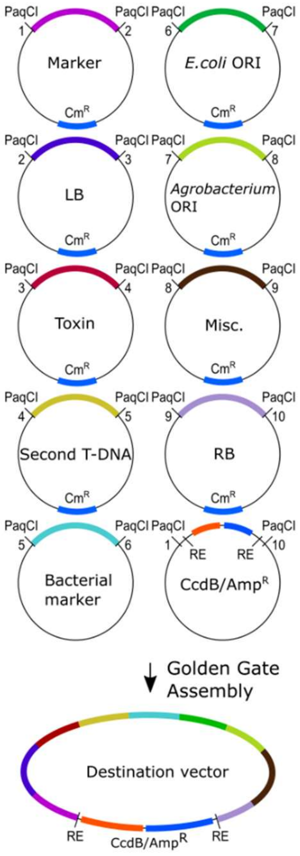


**Figure S1**: **The BackBone Builder (B3) cloning scheme.**

In the B3 cloning scheme, the Type IIS restriction enzyme PaqCI is used. Each entry plasmid part is flanked by PaqCI recognition sites with unique overhangs (1-10), that specify its position in the final destination vector. These entry plasmids contain individual backbone parts including an ORI, a bacterial selectable marker, and other required elements. There are ten entry plasmids labeled with number pairs, ranging from "1-2" to "9-10". A separate connector plasmid ("1–10") links the first and last positions and carries the negative selection marker ccdB together with a selectable antibiotic marker. This connector plasmid also contains additional sites (designated as restriction enzyme or "RE") that enable further cloning once the destination vector is assembled. These sites may be restriction enzyme sites, but they can also be non–restriction-based sequence elements such as homing endonuclease sites, loxP sites, or attL/attR sites, depending on the downstream cloning method. All entry plasmids share an identical backbone with a few exceptions. In positions "5-6" and "1-10", the bacterial marker is already part of the insert and therefore no chloramphenicol resistance marker is present in the backbone. In position "6-7", the pUC19 ORI is omitted in some entry plasmids to avoid incompatibility with the ORI being introduced. When a particular module is not required, a linker can be used instead. Following Golden Gate assembly and transformation in *E. coli*, selection is performed using $Amp^R$ and the antibiotic marker provided by entry plasmid "5-6". The resulting backbones can then serve as destination for inserting genes of interest using any preferred cloning system to generate expression vectors.

Every slot in the cloning system has a designated function (e.g., position 1–2 encodes a plant selectable marker gene). A brief explanation of the different categories is provided in the table below. Adhering to this scheme ensures compatibility with parts included in the kit as well as components generated by other researchers. Nevertheless, the structure can be adapted for research purposes, and the scheme can also be applied to other systems than *Agrobacterium* binary vectors, where alternative elements may be more appropriate.

**Table S1: Elements in the BackBone Builder (B3) cloning scheme.**

| Element | Function |
|---|---|
| Marker | Encodes a selectable or visual marker to enable selection of *Agrobacterium* transformed organisms. |
| LB | Left border of the T-DNA; terminates T-DNA transfer. |
| Toxin | Encodes a toxin located adjacent to the left border (LB), to select against organisms with backbone integration. |
| Second T-DNA | Independent T-DNA copy containing a selection marker that can integrate separately into the genome; enables isolation of marker-free transformants containing only the gene(s) of interest after segregation. |
| Bacterial marker | Selectable marker for bacterial transformation. |

| | |
|---|---|
| *E. coli* ORI | Origin of replication active in *E. coli*. Note that some origins function in both hosts. |
| *Agrobacterium* ORI | Origin of replication active in *Agrobacterium*. Note that some origins function in both hosts. |
| Misc. | Miscellaneous; may be used for various purposes, such as incorporating overdrive sequences, right border (RB) sequences, origins of transfer, replication proteins required for ORIs, or barcodes for experimental purposes. |
| RB | Right border of the T-DNA; initiates T-DNA transfer. |
| ccdB/AmpR | Cloning cassette containing ccdB for negative selection; other systems (e.g., sfGFP for visual screening) may be used instead. This part also contains the restriction sites or other sequences required for downstream cloning protocols. |

Empty entry vector cloning

The empty entry vectors are based on the amilCP_Orange chromoprotein vector (Addgene Plasmid #117850). First, we generated the entry vector for the "1-2" position by PCR amplifying with primer pairs 1-2 and 3-4 (primer sequences are listed in Supplementary Table S3). These PCR fragments were gel extracted and purified using the Zymoclean Gel DNA Recovery Kit (Zymo Research, D4001) and assembled with stitching oligos 5 and 6 according to De Saeger et al. (2022). The plasmid was extracted using the GeneJET Plasmid MiniPrep Kit (Thermo Fisher, K0503) according to the manufacturer's instructions.  The plasmid was digested with BglII (NEB, R0144S) according to the manufacturer's instructions. In a second round, PCR was initiated with primer pairs 7-8 and 9-10 using the templates "pGG-F-TaU3-ccdb-polyT-G" and "pGGB000". These PCR products were purified as described above and then assembled with the digested vector using NEBuilder® HiFi DNA Assembly Master Mix (NEB, E2621L) according to the manufacturer's instructions.

The other empty entry vectors were obtained by PCR and NEBuilder® HiFi DNA Assembly Master Mix (NEB, E2621L) using primers 11-26. The GGIB entry vector "10-1" was obtained by using the primer pairs 25-26 and 27-28 to amplify the plasmid amilCP_Orange chromoprotein vector mentioned above. PCR purification and assembly were done as described in the section above.

All vectors were verified by Whole Plasmid Sequencing (Eurofins).

Entry vector cloning

To construct new entry vectors for position n-n$^{+1}$, the entry vector n-n$^{+1}$ was digested with BsaI-HFv2 according to the manufacturer's instructions in a volume of 20 µL. PCR was used to amplify the part of interest using custom primers (Supplementary Table S3). PCR products were gel extracted using the Zymoclean Gel DNA Recovery Kit (Zymo Research, D4001). The products were then assembled into the digested vector using Gibson assembly according to De Saeger et al. (2022).

Alternatively, GreenGate parts (A-G) were used in a Golden Gate reaction to assemble the final vector according to Lampropoulos et al. (2013).

All plasmids were extracted using the GeneJET Plasmid MiniPrep Kit (Thermo Fisher, K0503) according to the manufacturer's instructions. Plasmids were extracted and sent for Sanger sequencing (Mix2Seq, Eurofins, Germany) using primers 159 and 160 (Supplementary Table S3). Parts that were too large to sequence with two Sanger reads were verified by Whole Plasmid Sequencing (Eurofins).

Golden Gate assembly

The reagents were mixed in a 0.2 mL PCR tube according to the reaction mix shown in the table on the left, and the reaction was run in a thermal cycler using the Golden Gate assembly program shown in the table on the right.

| **Entry** | **Volume (µL)** | **Amount** |
|---|---|---|
| 1-2 | 1 | 100 ng |
| 2-3 | 1 | 100 ng |
| 3-4 | 1 | 100 ng |
| 4-5 | 1 | 100 ng |
| 5-6 | 1 | 100 ng |
| 6-7 | 1 | 100 ng |
| 7-8 | 1 | 100 ng |
| 8-9 | 1 | 100 ng |
| 9-10 | 1 | 100 ng |
| 10-1 | 1 | 100 ng |
| T4 Ligase buffer (NEB, B0202S) | 1,5 | |
| T4 DNA Ligase (NEB, M0202L) | 0,75 | |
| PaqCI activator (NEB, R0745S) | 0,25 | |
| PaqCI (NEB, R0745S) | 0,75 | |
| $dH_2O$ | 2,75 | |
| *Total* | 15,00 | |

When certain positions were not needed, linkers were used instead. If several positions are spanned by one linker, 1 µL of linker was used and the rest was substituted with $dH_2O$.

**Golden Gate Assembly conditions**

| | | |
|---|---|---|
| 37°C | 15 min | |
| 37°C | 3 min | 50 X |
| 16°C | 3 min | |
| 37°C | 5 min | |
| 60°C | 5 min | |
| 16°C | ∞ | |

An additional PaqCI digestion step was included to remove assemblies that still contained intact PaqCI sites. For this, a 5× PaqCI master mix was prepared using the following composition:

| **Reagent** | **Volume (µL)** |
|---|---|
| rCutsmart buffer (NEB, B6004S) | 7,50 |
| PaqCI activator (NEB, R0745S) | 1,25 |
| PaqCI (NEB, R0745S) | 3,75 |
| *Total* | 12,50 |

A volume of 2.5 µL of this 5× mix was added to 12.5 µL of the assembly reaction and incubated at 37 °C for 1 hour. Subsequently, 2.5 µL of the digestion reaction was transformed into One Shot™ ccdB Survival™ 2 T1R Competent Cells (Thermo Fisher, A10460) according to the manufacturer's instructions. After incubation at 37°C for 1h, the cells were spun down at 1000 g and resuspended in 100 µL of SOC medium. This solution was plated on pre-warmed (37°C) LB medium containing the appropriate antibiotics. Single *E. coli* colonies were picked and grown overnight in 3 mL of LB medium containing the appropriate antibiotics. All plasmids were extracted using the GeneJET Plasmid MiniPrep Kit (Thermo Fisher, K0503) according to the manufacturer's instructions. Vectors were verified by restriction digestion and whole-plasmid sequencing (Eurofins). For restriction analysis, 5 µL of plasmid preparation was digested with XhoI (Promega, R6161), with a NotI–BglII double digest (Promega; NotI R6431, BglII R6621), or with a PvuI–XbaI double digest (Promega; PvuI R6321, XbaI R6181) in a total reaction volume of 10 µL, following the manufacturer's instructions. One plasmid from each construct was subsequently submitted for whole-plasmid sequencing (PlasmidSaurus).

Visual screening of colonies using eforCP marker

Each construct was revived from a glycerol stock by inoculating 5 mL of liquid medium (LB for *E. coli*, YEB for *Agrobacterium*), supplemented with the appropriate antibiotics. *E. coli* cultures were grown in LB with spectinomycin (100 µg/mL), and *Agrobacterium* cultures were grown in YEB with rifampicin (50 µg/mL), gentamicin (40 µg/mL), and spectinomycin (100 µg/mL). Cultures were incubated overnight at 37 °C (*E. coli*) or 28 °C (*Agrobacterium*). The next day, each culture was adjusted to an $OD_{600}$ of 0.1, and 40 µL was plated on LB or YEB agar (with the corresponding antibiotics). All 16 strains were plated on a single agar plate to allow direct comparison and imaging under identical conditions. An RGB image of the plate was recorded, and fluorescence images were acquired using a FastGene FAS-X imaging system (Nippon Genetics).

Leaf infiltration in *Nicotiana benthamiana*

*Agrobacterium tumefaciens* strains carrying the constructs of interest and the silencing suppressor P19 were grown overnight in 10 ml YEB medium supplemented with the appropriate antibiotics (gentamicin (40 µg/ml) + spectinomycin (100 µg/ml)) for assembly constructs; kanamycin (50 µg/ml) for P19) at 28 °C with agitation (180 rpm). The optical density at 600 nm ($OD_{600}$) of a 1:10 dilution was measured, and the culture volume required to obtain 2 ml at $OD_{600}$ = 1 was collected by centrifugation (10 min, 4000 rpm). Cell pellets were washed twice by resuspension in infiltration buffer, followed by centrifugation (10 min, 4000 rpm). The infiltration buffer consisted of 94 ml Milli-Q water, 1 ml 1 M $MgCl_2$, and 5 ml 200 mM MES, and was prepared fresh on the day of infiltration. After washing, pellets were resuspended in infiltration buffer containing acetosyringone (final concentration: 100 µM) and incubated for ≥ 2 h at 28 °C with shaking (180 rpm). $OD_{600}$ measurements were repeated, and cultures were mixed to obtain a final $OD_{600}$ of 0.25 for each construct in a final volume of 1 ml, including P19. Transient expression assays were performed on 6-week-old *Nicotiana benthamiana* plants. Agroinfiltration was carried out on the abaxial side of fully expanded leaves using a 1 ml syringe without needle. A small scratch was made with a sterile scalpel prior to infiltration, and ~100 µl of each culture mixture was infiltrated into discrete regions of the same leaf. A total of 16 construct combinations were infiltrated per leaf, and the procedure was repeated on four independent plants (one leaf per plant). Phenotypes were documented 5 and 6 days post-infiltration (5-6 dpi) by standard RGB imaging.

Maize transformation

To generate the expression vector, the elements A-pZm13-B and E-tZm13-LoxPsym-F were synthesized based on Zm00001eb421470 (Hamilton *et al.*, 1992) and F-TBS-G (Hily *et al*., 2009) and cloned in pTwist-Amp. These were combined with pGG-B-linker-C, pGG-C-GFP-D and pGG-D-linker-E in pB3G3HI-AG to yield pB3G3HI-pZm13-YFP-tZm13 (Supplementary Data S6). Maize transformation and digital PCR were done according to Vandeputte *et al*. (2024).

Use of Artificial Intelligence tools

The authors used ChatGPT (OpenAI, version GPT-5.2) for language editing and improvement of clarity in selected sections of the manuscript. No AI tools were used for data analysis or scientific interpretation. The authors reviewed and take full responsibility for all content.

**Table S2: BackBone Builder (B3) vectors.** Overview of all empty B3 entry plasmids and Agrobacterium parts in the B3 library, including ID, name, description, insert size (bp), resistance marker, and Vector Vault (VV) ID (https://vectorvault.vib.be/collection). Empty entry vectors (E1–E9) can be used to generate new parts. These vectors contain BsaI sites (A–G) according to the GreenGate standard (Lampropoulos et al., 2013), although new elements may also be introduced using Gibson assembly. Positions 1–10 represent the modular cloning slots used to assemble *Agrobacterium* destination vectors. *Agrobacterium* parts (A1–A43) included in the B3 system are listed as well.

Note: Abbreviations for antibiotic resistance markers: Amp, ampicillin; Cm, chloramphenicol; Erm, erythromycin; Gm, gentamicin; Kan, kanamycin; Spec, spectinomycin; Tet, tetracycline.

| ID | Name | Description | Insert size (bp) | Resistance | VV ID |
|---|---|---|---|---|---|
| **Empty entry vectors** | | | | | |
| E1 | B3-1-2 | Position 1-2: Plant marker | / | Amp/Cm | 26_27 |
| E2 | B3-2-3 | Position 2-3: LB | / | Amp/Cm | 26_28 |
| E3 | B3-3-4 | Position 3-4: Toxin/marker | / | Amp/Cm | 26_29 |
| E4 | B3-4-5 | Position 4-5: Second T-DNA | / | Amp/Cm | 26_30 |
| E5 | B3-5-6 | Position 5-6: Antibiotic resistance marker | / | Amp/Cm | 26_31 |
| E6 | B3-6-7 | Position 6-7: *E. coli* ORI | / | Amp/Cm | 26_32 |
| E7 | B3-7-8 | Position 7-8: *Agrobacterium* ORI | / | Amp/Cm | 26_33 |
| E8 | B3-8-9 | Position 8-9: Miscellaneous | / | Amp/Cm | 26_34 |
| E9 | B3-9-10 | Position 9-10: RB | / | Amp/Cm | 26_35 |
| **1 – Marker – 2** | | | | | |
| A1 | 1-p35S::bar::tNOS-2 | Encodes BASTA resistance | 1714 | Cm | 26_36 |
| A2 | 1-pNOS::bar::tNOS-2 | Encodes BASTA resistance | 1134 | Cm | 26_37 |
| A3 | 1-pNOS::hpt::tNOS-2 | Encodes hygromycin resistance | 1765 | Cm | 26_38 |
| A4 | 1-pOsAct-hpt(int)-t35S-2 | Encodes hygromycin resistance (intronized) | 2881 | Cm | 26_39 |
| A5 | 1-pNOS::nptII::tNOS-2 | Encodes kanamycin resistance | 1389 | Cm | 26_40 |
| A6 | 1-pOLE1-OLE1-mRuby3-tNOS-2 | Fluorescent seed selection in *Arabidopsis* | 3245 | Cm | 26_41 |
| A7 | 1-LINKER-2 | Linker | 24 | Cm | 26_42 |
| **2 – Left border – 3** | | | | | |
| A8 | 2-LB$^{pTiC58}$-3 | pTiC58 left border | 333 | Cm | 26_43 |
| A9 | 2-2xLB$^{pTiC58}$-3 | Double minimal pTiC58 left border | 75 | Cm | 26_44 |
| A10 | 2-LB$^{pTIT37}$-3.gb | pTIT37 left border | 310 | Cm | 26_45 |
| **3 – Toxin/maker – 4** | | | | | |

| | | | | | |
|---|---|---|---|---|---|
| A11 | 3-pNOS::Barnase$^{intron}$::tMAS-4 | Intronized barnase toxin driven by NOS promoter | 1058 | Cm | 26_46 |
| A12 | 3-LINKER-4 | Linker | 40 | Cm | 26_47 |
| **4 – Second T-DNA – 5** | | | | | |
| A13 | 4-RB-p35S::bar::t35S-LB-5 | Separate T-DNA cassette encoding BASTA resistance | 2425 | Cm | 26_48 |
| A14 | 4-RB-p35S::hpt(int)::t35S-LB-5 | Separate T-DNA cassette encoding hygromycin resistance | 2465 | Cm | 26_49 |
| A15 | 4-LINKER-5 | Linker | 40 | Cm | 26_50 |
| **5 – Antibiotic resistance marker – 6** | | | | | |
| A16 | 5-ermC-6 | Erythromycin resistance | 1060 | Erm | 26_51 |
| A17 | 5-aacC1-6 | Gentamycin resistance | 795 | Gm | 26_52 |
| A18 | 5-nptII-6 | Kanamycin resistance | 974 | Kan | 26_53 |
| A19 | 5-aadA-6 | Spectinomycin resistance | 1135 | Spec | 26_54 |
| A20 | 5-tetA-6 | Tetracycline resistance | 1433 | Tet | 26_55 |
| **6 – *E. coli* ORI – 7 *** | | | | | |
| A21 | 6-Phage P1-7 | Low copy number ORI | 5759 | Cm | 26_56 |
| A22 | 6-P15A-7 | Medium copy number ORI | 598 | Cm | 26_57 |
| A23 | 6-ColE1-7 | Medium copy number ORI | 614 | Cm | 26_58 |
| A24 | 6-ColE1-ROP-7 | Medium copy number ORI | 868 | Cm | 26_59 |
| A25 | 6-pUC-7 | High copy number ORI | 614 | Cm | 26_60 |
| **7 – *Agrobacterium* ORI – 8 *** | | | | | |
| A26 | 7-pRI-8 | Low copy number ORI; highly stable, derived from the Ri-plasmid | 4635 | Cm | 26_61 |
| A27 | 7-oriV-8 | Low copy number ORI; derived from RK2 plasmid; requires TrfA | 611 | Cm | 26_62 |
| A28 | 7-pSa-8 | Low copy number ORI; derived from pSa plasmid; requires pSa RepA | 452 | Cm | 26_63 |
| A29 | 7-pVS1+RepA-8 | Medium copy number ORI; broad-range origin of replication | 3020 | Cm | 26_64 |
| A30 | 7-WKS1-8 | Unknown copy number; derived from *Paracoccus* sp. plasmid | 1269 | Cm | 26_66 |
| **8 – Miscellaneous – 9** | | | | | |
| A31 | 8-trfA-9 | Replication initiation protein needed for oriV plasmids | 1319 | Cm | 26_67 |
| A32 | 8-pSa RepA (WT)-9 | Wild type pSa RepA | 1357 | Cm | 26_68 |
| A33 | 8-pSa RepA (K155N)-9 | Variant of pSa RepA with increased plant transformation efficiency (Szarzanowicz et al., 2025) | 1357 | Cm | 26_69 |

| | | | | | |
|---|---|---|---|---|---|
| A34 | 8-OriT-9 | Minimal origin of transfer | 127 | Cm | 26_70 |
| A35 | 8-LINKER-9 | Linker | 40 | Cm | 26_71 |
| **9 – Right border – 10** | | | | | |
| A36 | 9-minRB$^{pTiT37}$-10 | Minimal RB from pTiT37 | 191 | Cm | 26_72 |
| A37 | 9-RB$^{pTiT37}$-10 | RB from pTiT37 | 444 | Cm | 26_73 |
| **10 – Cloning cassette – 1** | | | | | |
| A38 | B3-U1-ccdB-U9 | Cassette to construct GGIB compatible vectors | 1512 | Cm | 26_74 |
| A39 | B3-A-ccdB-G | Cassette to construct GreenGate compatible vectors | 1443 | Cm | 26_75 |
| **Multi-slot-spanning modules** | | | | | |
| A40 | 1-LINKER-5 | Linker | 40 | Cm | 26_76 |
| A41 | 3-LINKER-5 | Linker | 40 | Cm | 26_77 |
| A42 | 7-LINKER-10 | Linker | 40 | Cm | 26_78 |

* Note on copy numbers: Actual plasmid copy number per cell varies substantially depending on factors such as insert size, host strain, and growth conditions. Due to population heterogeneity and the large variability reported in the literature - for example, pUC-based origins are frequently cited as reaching several hundred copies per cell (Anindyajati et al., 2016; Lee et al., 2006), whereas other studies measured mean values of ~61 copies per cell (Shao et al., 2021) and up to 15 copies per cell (Jahn et al., 2016) - we classify origins broadly into high, medium, and low copy-number categories rather than specifying absolute values. Here we follow the classification framework of Wang and Joffré (2025) and group origins into low (~1–5), medium (~5–50), and high (>50 copies per cell), acknowledging that published values for individual origins vary widely. For *Agrobacterium*, fewer studies are available than for *E. coli*, but recent work has reported copy-number values for several origins. For oriV, pSa, and pVS1 average copy numbers of 1.2, 4.5, and 9.5 copies per cell, respectively, have been observed (Szarzanowicz et al., 2024). Separate work has shown that the pRI origin supports highly stable plasmid maintenance in *Agrobacterium*, presumably due to the very low copy nature of this origin of replication (McBride & Summerfelt, 1990). WKS1 is a recent addition to the *Agrobacterium* toolbox and, to our knowledge, has not yet been characterized experimentally for copy number (Kocaoglan et al., 2024). Please also note that mutations in replication proteins associated with ORIs can affect copy numbers.

**Supplementary Data S1: High-Fidelity set of PaqCI overhangs**

To design a High-Fidelity set of overhangs to use with PaqCI, we made use of the NEBridge GetSet® Tool (https://ligasefidelity.neb.com/getset/run.cgi) with the "PaqCI, 1xT4 DNA Ligase buffer, 37-16 cycling" option selected. While several sets are possible, we selected one of the sets that showed a 100% estimated ligation fidelity (Figure 1).

| | AGTA | TACT | ACGA | TCGT | CGGA | TCCG | ATAA | TTAT | CTTA | TAAG | CGTC | GACG | AAGG | CCTT | GAGC | GCTC | GTCA | TGAC | GGAA | TTCC |
|---|---|---|---|---|---|---|---|---|---|---|---|---|---|---|---|---|---|---|---|---|
| AGTA | 0 | 232 | 0 | 0 | 0 | 0 | 0 | 0 | 0 | 0 | 0 | 0 | 0 | 0 | 0 | 0 | 0 | 0 | 0 | 0 |
| TACT | 232 | 0 | 0 | 0 | 0 | 0 | 0 | 0 | 0 | 0 | 0 | 0 | 0 | 0 | 0 | 0 | 0 | 0 | 0 | 0 |
| ACGA | 0 | 0 | 0 | 191 | 0 | 0 | 0 | 0 | 0 | 0 | 0 | 0 | 0 | 0 | 0 | 0 | 0 | 0 | 0 | 0 |
| TCGT | 0 | 0 | 191 | 0 | 0 | 0 | 0 | 0 | 0 | 0 | 0 | 0 | 0 | 0 | 0 | 0 | 0 | 0 | 0 | 0 |
| CGGA | 0 | 0 | 0 | 0 | 0 | 239 | 0 | 0 | 0 | 0 | 0 | 0 | 0 | 0 | 0 | 0 | 0 | 0 | 0 | 0 |
| TCCG | 0 | 0 | 0 | 0 | 239 | 0 | 0 | 0 | 0 | 0 | 0 | 0 | 0 | 0 | 0 | 0 | 0 | 0 | 0 | 0 |
| ATAA | 0 | 0 | 0 | 0 | 0 | 0 | 0 | 192 | 0 | 0 | 0 | 0 | 0 | 0 | 0 | 0 | 0 | 0 | 0 | 0 |
| TTAT | 0 | 0 | 0 | 0 | 0 | 0 | 192 | 0 | 0 | 0 | 0 | 0 | 0 | 0 | 0 | 0 | 0 | 0 | 0 | 0 |
| CTTA | 0 | 0 | 0 | 0 | 0 | 0 | 0 | 0 | 0 | 241 | 0 | 0 | 0 | 0 | 0 | 0 | 0 | 0 | 0 | 0 |
| TAAG | 0 | 0 | 0 | 0 | 0 | 0 | 0 | 0 | 241 | 0 | 0 | 0 | 0 | 0 | 0 | 0 | 0 | 0 | 0 | 0 |
| CGTC | 0 | 0 | 0 | 0 | 0 | 0 | 0 | 0 | 0 | 0 | 0 | 177 | 0 | 0 | 0 | 0 | 0 | 0 | 0 | 0 |
| GACG | 0 | 0 | 0 | 0 | 0 | 0 | 0 | 0 | 0 | 0 | 177 | 0 | 0 | 0 | 0 | 0 | 0 | 0 | 0 | 0 |
| AAGG | 0 | 0 | 0 | 0 | 0 | 0 | 0 | 0 | 0 | 0 | 0 | 0 | 0 | 219 | 0 | 0 | 0 | 0 | 0 | 0 |
| CCTT | 0 | 0 | 0 | 0 | 0 | 0 | 0 | 0 | 0 | 0 | 0 | 0 | 219 | 0 | 0 | 0 | 0 | 0 | 0 | 0 |
| GAGC | 0 | 0 | 0 | 0 | 0 | 0 | 0 | 0 | 0 | 0 | 0 | 0 | 0 | 0 | 0 | 166 | 0 | 0 | 0 | 0 |
| GCTC | 0 | 0 | 0 | 0 | 0 | 0 | 0 | 0 | 0 | 0 | 0 | 0 | 0 | 0 | 166 | 0 | 0 | 0 | 0 | 0 |
| GTCA | 0 | 0 | 0 | 0 | 0 | 0 | 0 | 0 | 0 | 0 | 0 | 0 | 0 | 0 | 0 | 0 | 0 | 168 | 0 | 0 |
| TGAC | 0 | 0 | 0 | 0 | 0 | 0 | 0 | 0 | 0 | 0 | 0 | 0 | 0 | 0 | 0 | 0 | 168 | 0 | 0 | 0 |
| GGAA | 0 | 0 | 0 | 0 | 0 | 0 | 0 | 0 | 0 | 0 | 0 | 0 | 0 | 0 | 0 | 0 | 0 | 0 | 0 | 278 |
| TTCC | 0 | 0 | 0 | 0 | 0 | 0 | 0 | 0 | 0 | 0 | 0 | 0 | 0 | 0 | 0 | 0 | 0 | 0 | 278 | 0 |

| **Position** | **OH1-part-OH2** |
|---|---|
| 1-2 | 5'-AGTA-part-ACGA-3' |
| 2-3 | 5'-ACGA-part-CGGA-3' |
| 3-4 | 5'-CGGA-part-ATAA-3' |
| 4-5 | 5'-ATAA-part-CTTA-3' |
| 5-6 | 5'-CTTA-part-CGTC-3' |
| 6-7 | 5'-CGTC-part-AAGG-3' |
| 7-8 | 5'-AAGG-part-GAGC-3' |
| 8-9 | 5'-GAGC-part-GTCA-3' |
| 9-10 | 5'-GTCA-part-GGAA-3' |
| 1-10 | 5'-AGTA-part-GGAA-3' |

**Overhang design of B3. Left:** Ligation frequency matrix showing normalized ligation efficiency estimations. Dark blue denotes high Watson-Crick pairing probabilities. No undesired ligations are predicted using this overhang set**. Right:** Selected overhangs with their position in the B3 system.

**Supplementary Data S2: B3 cloning results**

**Table S5: Overview of B3 assemblies and corresponding expected restriction digestion patterns shown in Figure S5.** This table links each construct to its origin-of-replication combination, the restriction enzymes used for validation, and the expected fragment sizes. Sample IDs refer to the clone numbers prepared for analysis. Sets labeled “n” use an alternative replicate numbering format; for example, “5-n” corresponds to samples 5-1, 5-2, 5-3, 5-4, and 5-5.

| Assembly | ORI combination | Restriction enzyme | Fragment sizes (bp) | Samples |
|---|---|---|---|---|
| AG1 | pUC-pVS1 | PvuI + XbaI | 4151 + 2205 + 1458 + 679 | 1-5 |
| AG2 | ColE1-pVS1 | PvuI + XbaI | 4151 + 2205 + 1458 + 679 | 6-10 |
| AG3 | P15A-pVS1 | PvuI + XbaI | 4135 + 2205 + 1458 + 679 | 11-15 |
| AG4 | Phage P1-pVS1 | PvuI + XbaI | 5210 + 4086 + 2205 + 1458 + 679 | 16-20 |
| AG5 | pUC-pRI | XhoI | 4820 + 2512 + 1693 + 1084 | 5-n |
| AG6 | ColE1-pRI | XhoI | 4820 + 2512 + 1693 + 1084 | 6-n |
| AG7 | p15A-pRI | XhoI | 4820 + 2512 + 1693 + 1068 | 7-n |
| AG8 | Phage P1-pRI | NotI + BglII | 6424 + 4309 + 2731 + 1790 | 8-n |
| AG9 | pUC-WKS1 | PvuI + XbaI | 2401 + 2205 + 1458 + 679 | 41-45 |
| AG10 | ColE1-WKS1 | PvuI + XbaI | 2401 + 2205 + 1458 + 679 | 46-50 |
| AG11 | P15A-WKS1 | PvuI + XbaI | 2385 + 2205 + 1458 + 679 | 51-55 |
| AG12 | Phage P1-WKS1 | PvuI + XbaI | 4086 + 3460 + 2205 + 1458 + 679 | 56-60 |
| AG13 | pUC-OriV | PvuI + XbaI | 5227 + 1458 + 679 | 61-65 |
| AG14 | ColE1-OriV | PvuI + XbaI | 5227 + 1458 + 679 | 66-70 |
| AG15 | P15A-OriV | PvuI + XbaI | 5211 + 1458 + 679 | 71-75 |
| AG16 | Phage P1-OriV | PvuI + XbaI | 6286 + 4086 + 1458 + 679 | 76-80 |

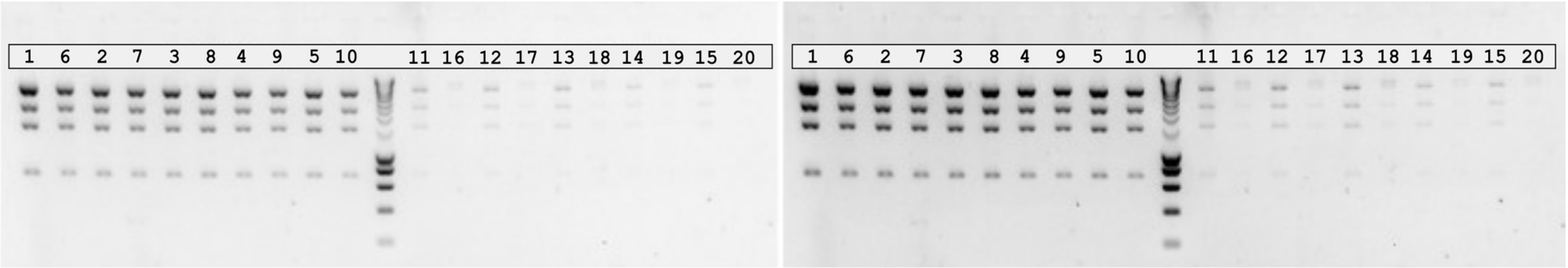


**Figure S2: Expected digestion patterns for AG1 (1-5), AG2 (6-10), AG3 (11-15), and AG4 (16-20).** Left: standard exposure; right: increased intensity of the same gel. Bands were compared against the BenchTop 1 kb DNA Ladder (Promega, G7541).

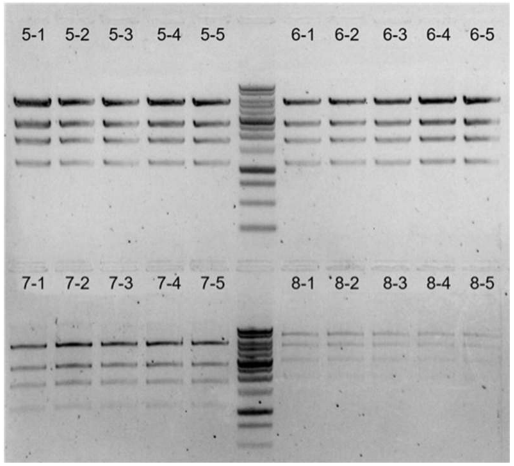


**Figure S3: Expected digestion pattern for AG5, AG6, AG7, and AG8 (sample 5-n, 6-n, 7-n, and 8-n, respectively).** Bands were compared against the BenchTop 1 kb DNA Ladder (Promega, G7541).

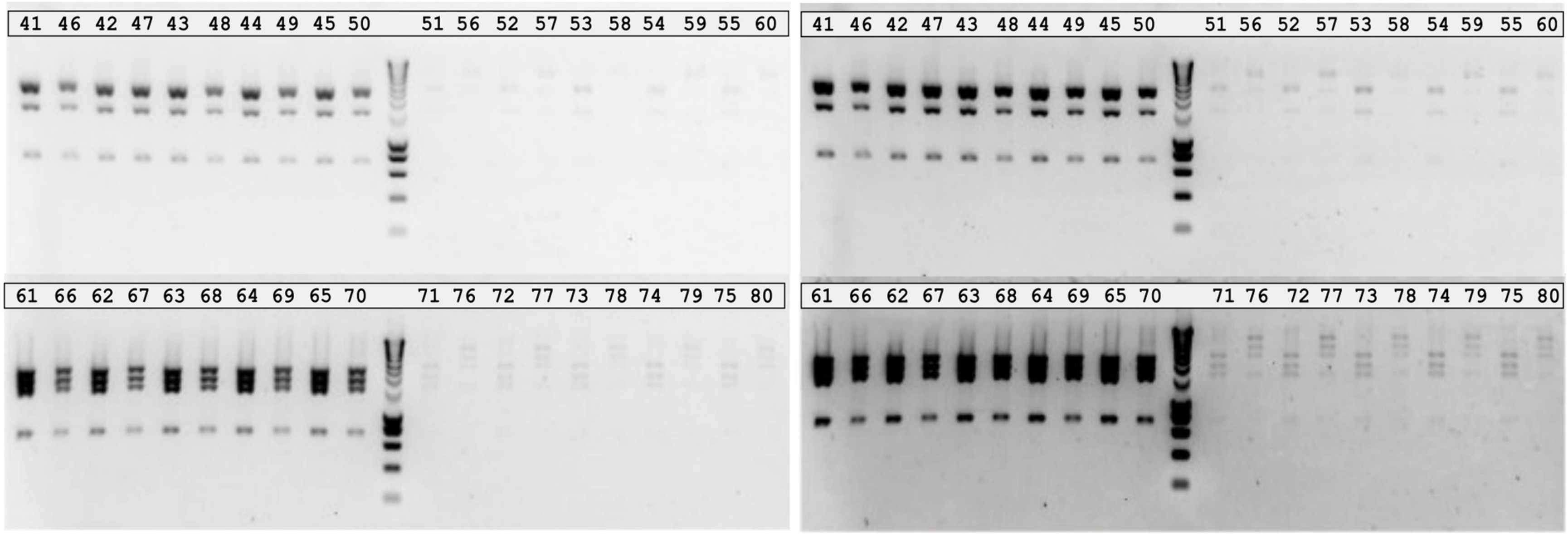


**Figure S4: Expected digestion patterns for AG9 (1-5), AG10 (6-10), AG11 (11-15), AG12 (16-20), AG13 (61-65), AG14 (66-70), AG15 (71-75) and AG16 (76-80).** Left: standard exposure; right: increased intensity of the same gel. Bands were compared against the BenchTop 1 kb DNA Ladder (Promega, G7541).

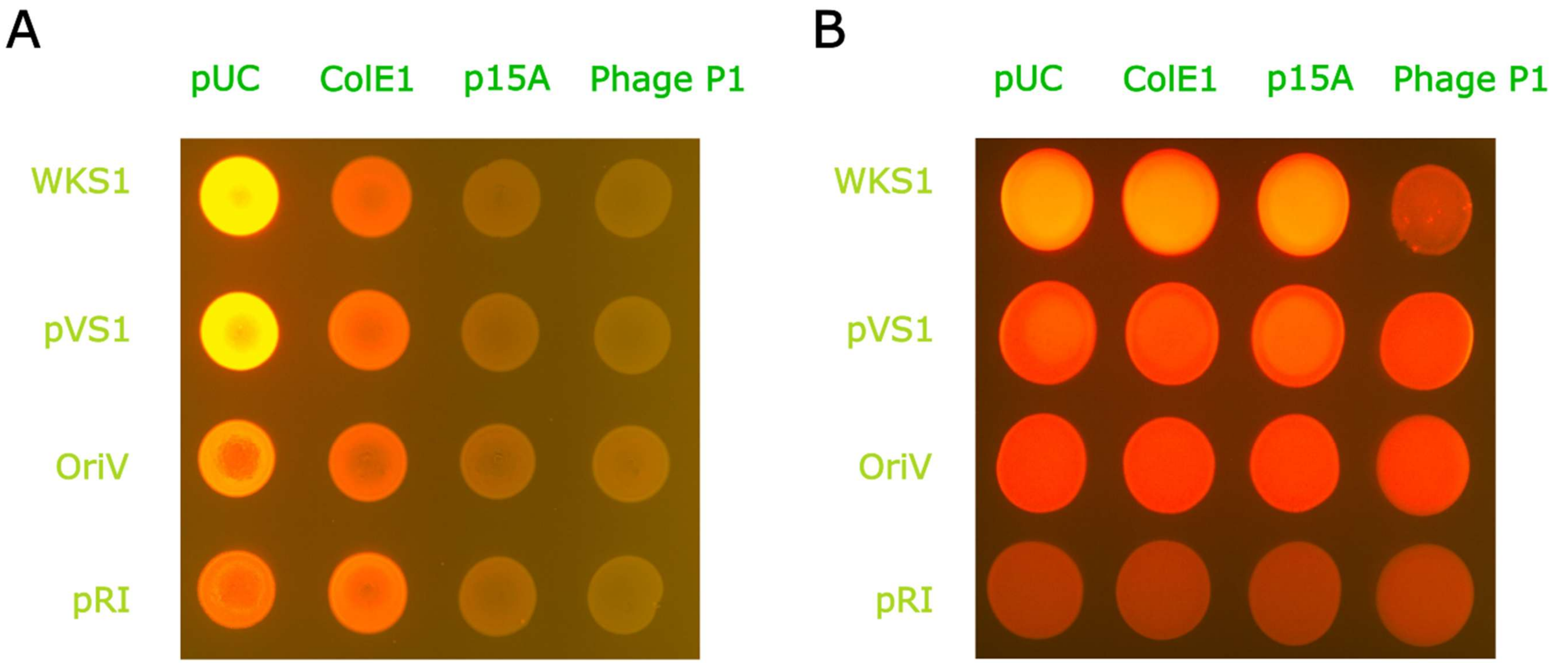


**Figure S5: Fluorescence images of *E. coli* and *A. tumefaciens* carrying the 16 ORI-matrix constructs. (A)** *E. coli* DH5α transformed with the 16 ORI-matrix vectors. **(B)** *Agrobacterium tumefaciens* LBA4404 transformed with the same 16 ORI-matrix vectors. Images correspond to the *same* physical plates as those shown in Figure 2B–C.

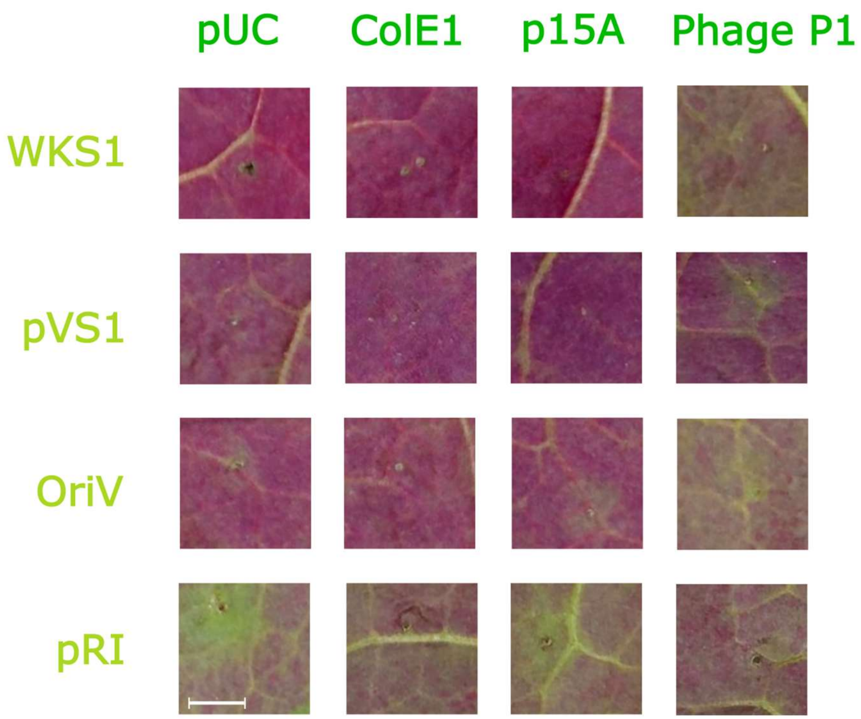


**Figure S6: Transient RUBY accumulation in *Nicotiana benthamiana* leaves infiltrated with 16 ORI combinations, imaged at 6 days post-infiltration (6 dpi).** Scale bar = 5 mm.

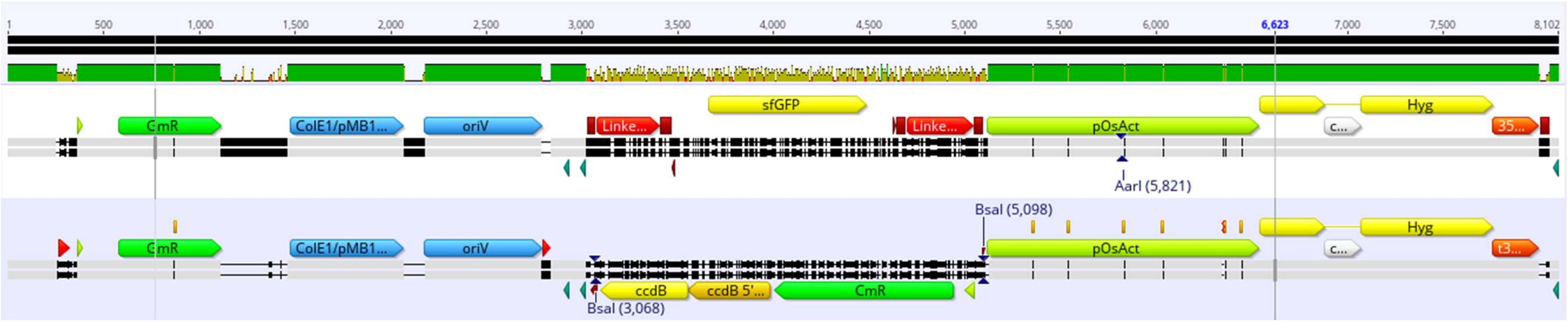


**Figure S7: Comparison of pG3HI-AG (upper) and pB3G3HI-AG (lower) plasmids.** Pairwise sequence alignment visualization showing nucleotide identity across the two constructs. The discontinuity around 3–5 kb corresponds to the cloning cassette, which differs between the plasmids (sfGFP in pG3HI-AG versus ccdB in pB3G3HI-AG).
